# Molecular Basis for Reduced Cleavage Activity and Drug Resistance in D30N HIV-1 protease

**DOI:** 10.1101/2021.02.28.433284

**Authors:** Subhash C Bihani, Madhusoodan V Hosur

**Affiliations:** Protein Crystallography Section, Radiation Biology & Health Sciences Division, Bhabha Atomic Research Centre, Trombay, Mumbai 400085, India; Homi Bhabha National Institute, Anushaktinagar, Mumbai 400094, India; National Institute of Advanced Studies, IISc campus, Bengaluru 560012, India

## Abstract

Nelfinavir is one of the FDA approved HIV-1 protease inhibitors and is a part of HAART therapy for the treatment of HIV-AIDS. Nelfinavir was the first HIV-1 protease inhibitor to be approved as a Paediatric formulation. The application of HAART had resulted into significant improvement in the life of AIDS patients. However, emergence of drug resistance in HIV-1 protease limited the use of many of these drugs including nelfinavir. A unique mutation observed frequently in patients treated with nelfinavir is D30N as it is selected exclusively by nelfinavir. It imparts very high resistance to nelfinavir but unlike other primary mutations does not give cross resistance to the majority of other drugs. D30N mutation also significantly reduces cleavage activity of HIV-1 protease and affects the viral fitness. Here, we have determined structures of D30N HIV-1 protease in unliganded form and in complex with the drug nelfinavir. These structures provide rationale for reduced cleavage activity and molecular basis of resistance induced by D30N mutation. The loss of coulombic interaction part of a crucial hydrogen bond between the drug and the enzyme, is a likely explanation for reduced affinity and drug resistance towards nelfinavir. The decreased catalytic activity of D30N HIV protease, due to altered interaction with substrates and reduced stability of folding core may be the reasons for reduced replicative capacity of the HIV harboring D30N HIV-1 protease.

## Introduction

Development of HIV-1 protease inhibitors is a successful story of structure-based drug design approach. Introduction of these inhibitors as part of HAART therapy has improved the life of millions of patients. Together with the HIV-1 prevention programs, these drugs have successfully reduced the burden of HIV-1 AIDS on the human population. So far, 8 HIV protease inhibitors have been approved by food and drug administration, USA. All these drugs are competitive inhibitors *i.e.,* their effectiveness is based on balance between their affinity versus that of the substrates in the binding site. Drug resistance mutations against all the approved protease inhibitors have been reported. Many of these drug resistance mutations are known to affect the catalytic efficiency of the enzyme but the effect is more on the inhibitor binding as compared to the substrate. Thus under the selection pressure of these drugs, mutated protease proves advantageous despite altered catalytic proficiency. Also, accessory mutations restores the catalytic activity of the enzyme providing more selective advantage to mutant protease.

Nelfinavir mesylate, Viracept is one of the FDA approved HIV protease inhibitors developed by Agouron Pharmaceuticals using structure based drug design strategy and Monte Carlo simulations [Gehlhaar et al 1995]. Recently, nelfinavir is also shown to be effective against Covid-19 as it inhibits SARS-CoV-2 3CL protease (Jan et al., 2021). Nelfinavir has also shown anti-cancer properties through its effects on different cellular conditions like cell cycle, apoptosis, autophagy, the proteasome pathway, oxidative stress, the tumor microenvironment, and multidrug efflux pumps (Subeha and Telleria, 2020). Multiple clinical trials have supported the anticancer applications of nelfinavir with one of the clinical trials showing it as a promising drug for long-term local control and survival in patients with locally advanced non–small cell lung cancer (Rengan et al., 2019). Nelfinavir has a S-phenyl group at P1 subsite and a novel 2-methyl-3-hydroxy benzamide group at P2 subsite along with a dodecahydroisoquinoline ring at P1’ site and tert-butylcarboxamide group at P2’ site (Figure 1). Nelfinavir has high affinity towards HIV-1 protease with Ki of 2 nM and was the first protease inhibitor approved for use in pediatric AIDS patients. Nelfinavir has a unique drug resistance mutation pattern in comparison with other FDA approved protease inhibitors. D30N, L90M, and N88S are the major mutations observed against nelfinavir along with many minor mutations including M36I, M46I/L, I54V, A71V, V82A, I84V, and N88D *etc* [Rhee et al 2003]. These mutations render the drug ineffective and require continuous development of more effective HIV-1 protease inhibitors which needs a clear understanding of the structural effects caused by these drug resistance mutations. For present work, mutations D30N in which an active site aspartate is replaced with asparagine is chosen and their crystal structures both in unliganded form and in complex with the drug nelfinavir were determined. D30N is the most common mutation observed in the 36-59% population of HIV infected patients undergoing nelfinavir therapy [Patick et al 1998, Yerly et al 2001, Perrin and Mammano 2003, Saah et al 2003]. Interestingly, D30N is exclusively selected by nelfinavir and does not provide cross resistance to the majority of other HIV protease inhibitors [Perrin and Mammano 2003]. D30N mutation leads to ~39 fold resistance against nelfinavir as measured by PhenoSense assay [Rhee et al 2003], while the increase in K_i_ value of approximately 20-fold compared to the wild type is reported [Kozisek et al 2007]. As compared to other HIV protease drug resistance mutations, D30N also has a crippling effect on the catalytic activity of HIV-1 protease with upto 90% loss in the k_cat_ value [Mahalingam et al 1999, Kozisek et al 2007]. Phage λ based genetic screening method has shown 97.8% loss of proteolytic activity in D30N HIV-1 protease as compared to the wild type HIV-1 protease [Parera et al 2007]. Similarly, highly deleterious effect of D30N on the virus and its replicative capacity has been observed [Martinez-Picado et al 1999]. Earlier a low-resolution structure of D30N HIV-1 protease-nelfinavir complex suggested that reduced binding affinity of nelfinavir is not due to structural changes in the active site pockets [Kozisek et al 2007]. High-resolution crystal structure of D30N mutant HIV-1 protease will help in understanding the structural effects of D30N, its molecular basis of drug resistance and exclusivity against nelfinavir. Also, there is no structure available for D30N HIV-1 protease in unliganded form, which is important to completely understand the structural effects caused by mutations. This paper reports X-ray crystal structures of D30N HIV-1 protease both in unliganded form and in complex with the drug nelfinavir determined to resolutions of 1.65 Å and 1.91 Å respectively. These structures will provide atomic level details of the structural effects caused by the mutation D30N and also reveal the molecular basis of drug resistance. The information gained will be highly beneficial for modification of nelfinavir to keep it relevant and development of new HIV-1 protease inhibitors.

**Figure 1:**
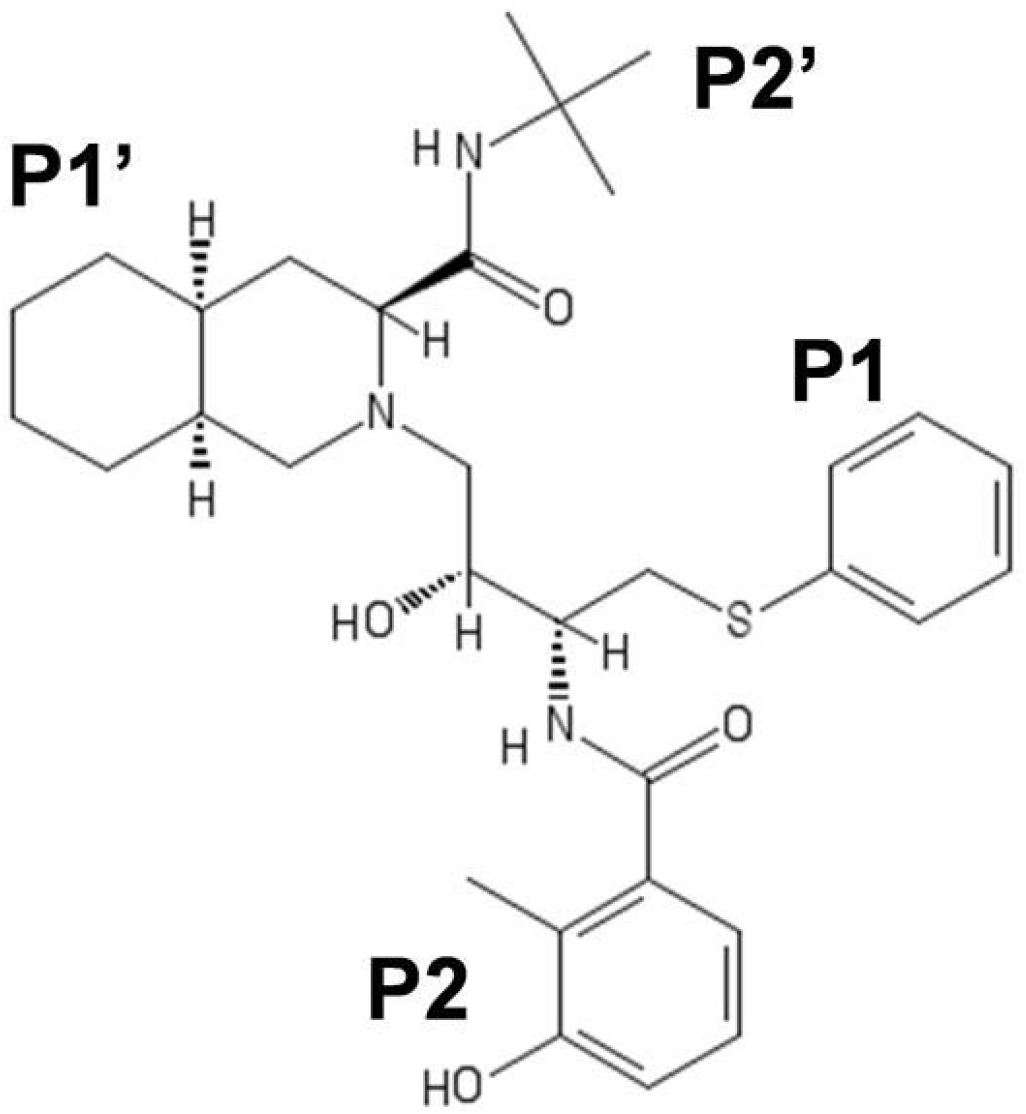
A 2-D schematic diagram of nelfinavir.

## Material and methods

### Site-directed mutagenesis, purification, and crystallization of D30N HIV-1 protease

The tethered dimer of HIV-1 protease gene described earlier was used as a template to create D30N HIV-1 protease (Bihani et al., 2009; Prashar et al., 2010). Since HIV-1 protease is active as a homodimer, two HIV-1 protease genes are fused with a GGSSG linker to express it as a tethered dimer. Since there are two chains, mutation has to be introduced in both the chains, hence, the mutations were introduced by Quick-change multisite directed mutagenesis method [Stratagene, La Jolla, CA]. The primer sequence used was 5’-CTGGATACCGGTGCTGAT*A*ATACTGTACTGGAGGAG -3’ and was designed using Primer X software [Lapid 2003]. Presence of mutations in the protease gene was confirmed by DNA sequencing and D30N HIV-1 protease gene insert was transformed into *E.coli* BL21 (DE3) cells. The cells were grown in Luria-Bertani (LB) broth till OD_600_ of 0.6-0.8 and protein expression was induced by adding 1 mM IPTG. Cells were harvested after 3-4 hours by centrifugation at 5000 x g for 10 minutes at 4°C. Temperature for all the subsequent steps was maintained at 4°C and all the buffers were pre-cooled to 4°C. The cell pellet were lysed by ultrasonication in lysis buffer (20mM Tris, 10mM EDTA and 1% Triton X-100) in pulse mode (1 sec on and 2 sec off- total 4 minutes pulse on). The lysate was then centrifuged at 10000 x g for 10 minutes. D30N HIV-1 protease accumulated in inclusion bodies, hence, the supernatant was discarded and grayish white pellet of inclusion bodies was retained. The pellet was washed thrice with lysis buffer using ultrasonication and centrifugation to remove all the impurities. The washed pellet which mostly contains pure D30N HIV-1 protease was finally dissolved in 3-4 ml of 67% acetic acid followed by centrifugation at 10000 x g for 15 min. The supernatant was diluted by 33 times with distilled water and dialyzed twice against cold distilled water and finally against ice-cold refolding buffer (20 mM PIPES pH 6.5, 100mM NaCl, 1mM DTT and 10% glycerol). After removing the precipitate by centrifugation at 10000x g for 15 minutes, the purified protein was concentrated by centrifugal concentrators up to the desired concentration. The final protein concentration for crystallization trials was 1-3 mg/ml in 50mM sodium acetate buffer (pH 4.5) containing 1mM DTT. Crystallization was done at room temperature by the hanging-drop vapor diffusion method. Hexagonal rod shaped crystals were obtained with sodium citrate/ sodium dihydrogen phosphate as reservoir buffer (200/100mM, pH 6.2) and ammonium sulfate (1-6% of saturated solution) as precipitant. Mutant HIV-1 proteases were also co-crystallized with the drug nelfinavir. Five molar excess of nelfinavir mesylate (received as gift from Ms. Cipla, India) dissolved in 100% DMSO was mixed with D30N HIV-1 protease and incubated at room temperature for 2-3 hours before setting up the crystallization experiments. Crystals of D30N HIV-1 protease and nelfinavir complex were obtained in conditions similar to that of unliganded D30N HIV-1 protease.

### Diffraction data collection

X-ray diffraction data were collected at ESRF, Grenoble, France, by the oscillation method, under cryo conditions using a crystal equilibrated in cryoprotectant solution (25% glycerol and 75% reservoir solution). For the D30N unliganded crystal, 90 frames, each for an oscillation angle of 1° were collected. For the D30N-nelfinavir complex crystal, 70 frames, each for an oscillation angle of 1° were collected. The oscillation frames were indexed, integrated and scaled with program suite XDS [Kabsch 2010]. Data collection statistics for both the crystals are summarized in Table 1.

**Table 1:** Data collection statistics of D30N unliganded and D30N-nelfinavir structures

| **Data-collection statistics** | **D30N** | **D30N-Nelf** |
| --- | --- | --- |
| Wavelength (Å) | 0.934000 | 0.979778 |
| Detector distance (mm) | 162 | 200 |
| Space group | P6<sub>1</sub> | P6<sub>1</sub> |
| Unit-cell parameters (Å, °) | 62.37, 62.37, 82.3<br>90, 90, 120 | 62.05, 62.05, 81.46<br>90, 90, 120 |
| Mosaicity (°) | 0.131 | 0.344 |
| Resolution range (Å) | 45.16 - 1.651<br>(1.75-1.65) | 26.87 - 1.91<br>(2.01-1.91) |
| Total No. of reflections | 121767(18693) | 58940 (8236) |
| No. of unique reflections | 21751 (3438) | 13778 (1951) |
| Completeness (%) | 99.5 (97.8) | 99.5 (99.2) |
| Mean I/σ(I) | 24.2 (3.1) | 12.5 (3.1) |
| R<sub>merge</sub>(%) | 4.4 (58.1) | 7.7 (49.1) |
| **Structure Refinement** |  |  |
| Rwork (%) | 19.1 | 18.2 |
| Rfree(%) | 22.5 | 22.9 |
| RMSD bond | 0.006 | 0.007 |
| RMSD angle | 0.88 | 0.995 |
| Ramchandran plot, favored | 100.00 | 98.4 |
| Ramchandran plot, allowed | 0.00 | 1.6 |
| PDB Code | 7DPQ | 7DOZ |

### Structure refinement

D30N HIV-1 protease and D30N HIV-1 protease - nelfinavir complex structures were determined by molecular replacement as implemented in PHENIX software suite. Structure of N88D HIV-1 protease tethered dimer (PDB Id 3KT2) was used as the phasing model. The structures were refined using crystallography software suite PHENIX [Liebschner et al., 2019]. In the intial stage, a rigid body refinement and simulated annealing step followed by restrained refinement was performed. Only restrained refinement without simulated annealing was used at all subsequent stages. Translational/Liberation/Screw (TLS) refinement as implemented in the PHENIX was performed at final stages [Painter and Merritt 2006]. Water molecules were added automatically by the PHENIX and validated manually along with model building using WinCoot [Emsley and Cowtan 2004, Emsley et al 2010]. Figures were prepared using the software PyMol [Schrodinger, LLC].

## Results and discussion

The D30N HIV-1 protease unliganded and in complex with nelfinavir were crystallized in P6_1_ space group with one tethered dimer in an asymmetric unit. The residues in the first monomer are numbered as 1–99 and those in the second monomer are numbered 1001–1099. Linker region of the tethered dimer is not modeled in the present structures as there is no electron density visible for it suggesting disordered nature of this region. Crystals of D30N HIV-1 protease and D30N-nelfinavir complex crystals diffracted to resolution of 1.65 Å and 1.91 Å respectively and structures were refined to R_free_ values of 22.5% and 22.9% respectively with very good stereochemistry. Refinement statistics for both the structures are given in Table 1.

### Nelfinavir is bound in two orientations

The clear difference density visible in the active site of D30N-nelfinavir complex structure displayed an approximate two-fold symmetry suggesting that nelfinavir is bound in two orientations corresponding to the pseudo-two fold symmetry of the active site. Therefore, nelfinavir is modeled with two alternate conformations (A and B) (Figure 2). The two orientations A and B were included with occupancy of 0.5 each at the start of the refinement which converged to the occupancy of 0.49/0.5 respectively at the end of refinement.

**Figure 2:**
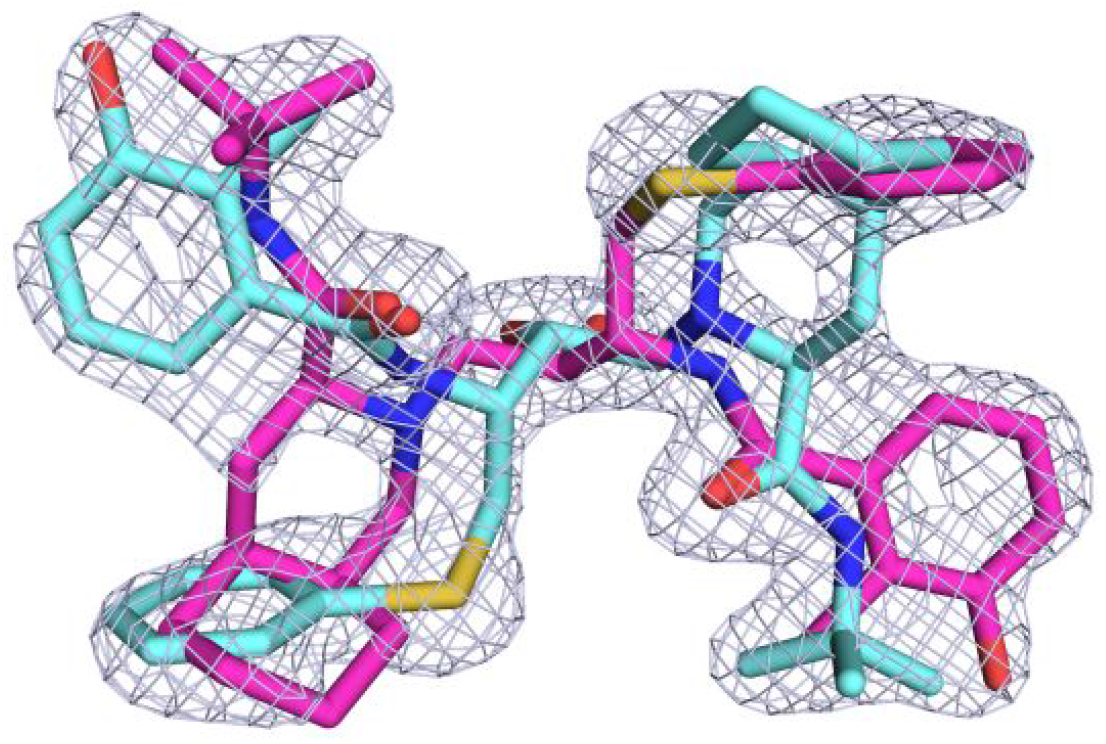
Two fold orientation of nelfinavir and 2mFo-dFc map at 1σ level. Orientation A is shown in magenta and B in cyan.

### Orientation of side chain of asparagine 30

Simulated annealed OMIT density calculated by omitting either the whole amino acid residue N30/1030 or only the ND2 atom in the side chain, clearly show that the positions of the 30/1030 ND2/OD1 atom in the present structure have been accurately determined (Figure 3). However, Asn30 side chain amide group in the D30N HIV-1 protease active site can take two different orientations. Since, current resolution of the X-ray structures do not permit identification of H atoms on NH_2_, and distinguishing between N and O atoms based on electron density map is not possible, assigning correct orientation of Asn30 side chain is difficult. Hydrogen bonding interactions and rotamer preference can help to overcome this problem. However, Asn30 amide group can acts both as donor and acceptor depending upon its orientation. Also, hydroxyl O of 2-methyl-3-hydroxyl benzamide group in nelfinavir can acts both as an hydrogen bond acceptor or donor. In such scenarios, where the interacting atoms are both potential donors and acceptors, further analysis of the surrounding environment can help in correct assignment [Word et al 1999]. Therefore, in D30N-nelfinavir complex structure, both the scenarios of Asn30 in alternate orientations (whether ND2 or OD1 of Asn30 is towards the nelfinavir) were compared. Both the situations resulted into very similar R-factors during refinement. Analysis by MolProbity server also suggested that both the situations are equally likely [Christopher et al 2018]. Rotamer with ND2 towards nelfinavir is favored (47%) over rotamer where OD2 is towards nelfinavir (4%) configuration in standard rotamer library for asparagine in WinCoot. Hence in the final structure, we have kept the orientation where ND2 of Asn30 is towards the nelfinavir.

**Figure 3:**
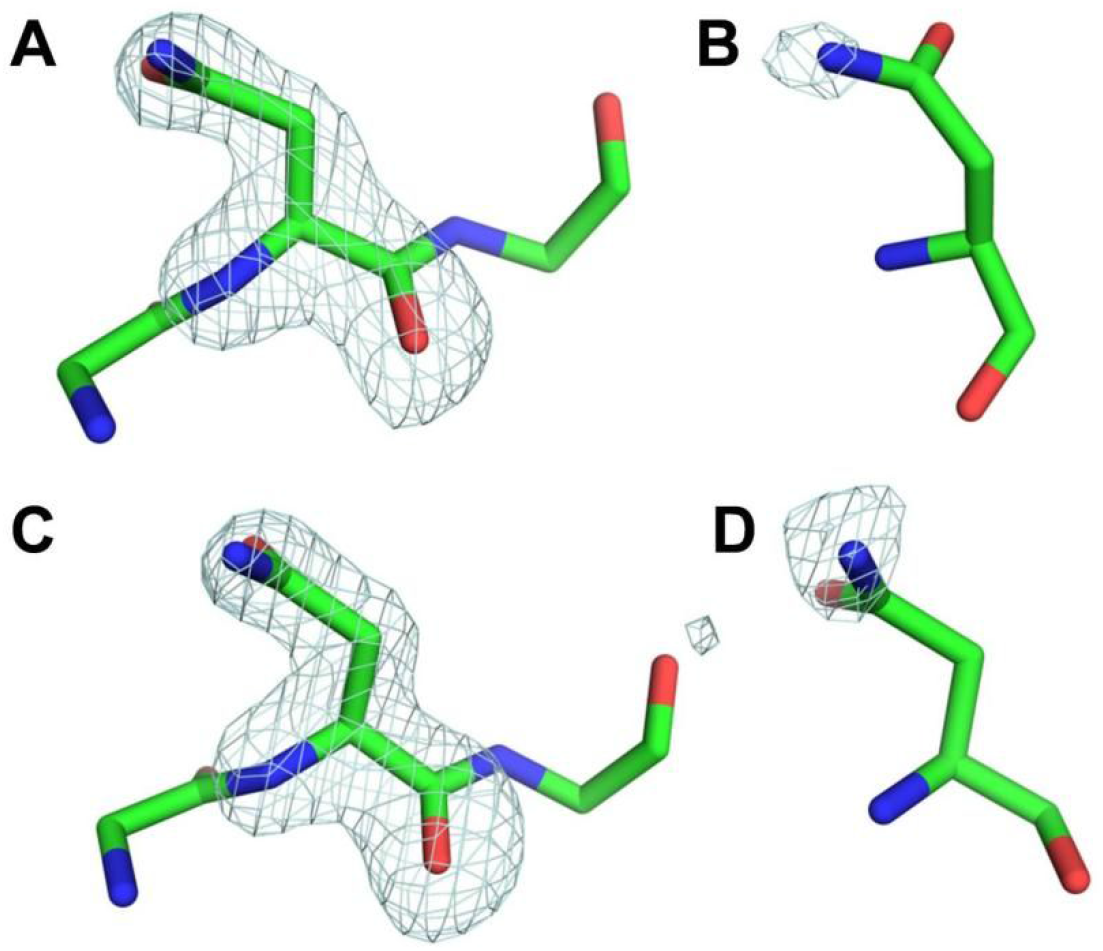
Simulated annealed omit map of D30N HIV-1 protease -nelfinavir complex structure (PDB ID 7DOZ) at 3σ level for (A) N30 (B) N30 ND2 (C) N1030 and (D) N1030 ND2

### Comparing molecular conformation of D30N HIV-1 protease with wild-type HIV-1 protease

The D30N unliganded structure superposed well on the unliganded tethered HIV-1 protease structure [PDB ID 3G6L] with a RMSD of 0.53 Å. Overall conformation is similar for both the structures with deviations at D25 Cα positions being less than 0.1Å (0.05/0.07Å), suggesting that D30N mutation does not have any major structural effect. All the residues that deviated by more than 0.5 Å were found to be on the surface and in the loop regions, which are intrinsically flexible, and have been found to have higher RMSDs in different HIV-1 protease structures. However, at the mutation site, deviations at Cα positions of 30^th^ residues are (0.16Å/0.16Å) with much higher deviations (0.69Å/0.49Å) observed at side chain residues measured from OD2 to ND2 position as N30 in D30N moves outward towards the residue N88.

### Comparing molecular conformation of D30N protease-nelfinavir complex with wild-type protease-nelfinavir complex

Despite having been crystallized in different space groups, D30N-nelfinavir complex structure superposes on wild type-nelfinavir complex structure (PDB ID 3EKX) with RMSD of 0.427Å, suggesting that overall conformation remain same with deviations at D25 Cα positions being less than 0.1Å (0.08/0.05Å). This RMSD of 0.427Å is even less than 0.53Å RMSD between unliganded structures, which were crystallized in the same space groups. This suggests that there are some differences in the wild type and mutant proteases and those were reduced upon binding to nelfinavir. Interestingly, deviations at D/N30 Cα is 0.05/0.08Å and at side chain ND2/OD2 atoms is 0.17/0.34Å, which is much less than the deviation observed in unliganded structures. Much larger shifts of the order of 0.7 Å– 0.8 Å were predicted for these residues by theoretical simulations [Ode et al 2007]. Reduced deviations in the positions of 30^th^ residues in complex structures suggest presence of induced fit due to the binding of nelfinavir. Overall these analyses suggest a preconfigured active site in the wild type protease that accepts nelfinavir without much change while in D30N HIV-1 protease small conformational changes are required to fit nelfinavir. Nelfinavir binds to the active site of both mutant and wild type protease in similar orientation and conformation with small displacements distributed in all four subsites (Figure 4).

**Figure 4.**
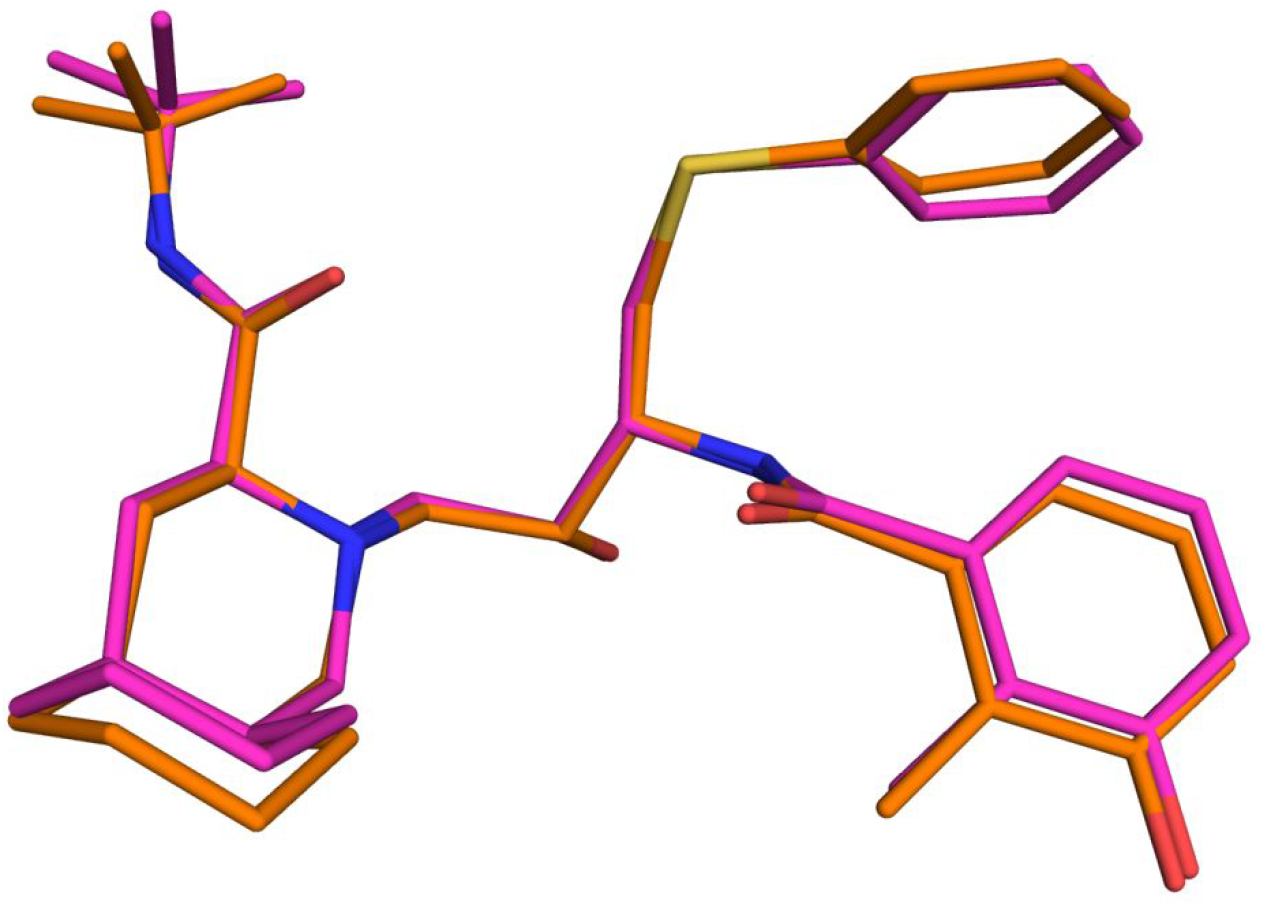
Superposition of nelfinavir in D30N-nelfinavir complex (magenta) with wild-type-nelfinavir complex structure (green).

### Comparing hydrogen bonding interactions in D30N-nelfinavir complex with wild type-nelfinavir complex structure

The hydrogen bonding interactions (donor-acceptor distance less than 3.2Å) of nelfinavir in the binding cavity of D30N HIV-1 protease are shown in Figure 5. Nelfinavir molecule has four oxygen in two hydroxyl and two carbonyl groups, all of which form hydrogen bonds, either directly or through a water molecule. Similar to wild type protease-nelfinavir complex, the central secondary hydroxyl group (O21) of nelfinavir is at distances of 2.6Å- 3.0Å from all four oxygens of the catalytic aspartates in D30N HIV-1 protease-nelfinavir complex. A bridging water known as flap water (FW) mediates hydrogen bonds between carbonyl oxygens O17 and O25 of the drug and I50 and I1050 main chain amide nitrogen atoms of D30N HIV-1 protease. Even at the mutation site, the N30 side chain retains the hydrogen bond (d = 2.8 Å) with hydroxyl oxygen O38 of 2-methyl-3-hydroxy benzamide group from nelfinavir (Figure 5). Similar to the wild type HIV-1 protease, water mediated hydrogen bond connects N12 of nelfinavir with residue D1029. These interactions, along with the hydrophobic interactions are probably responsible for the proper positioning of the tert-butyl group in the S2’ subsite of HIV-1 protease.

**Figure 5:**
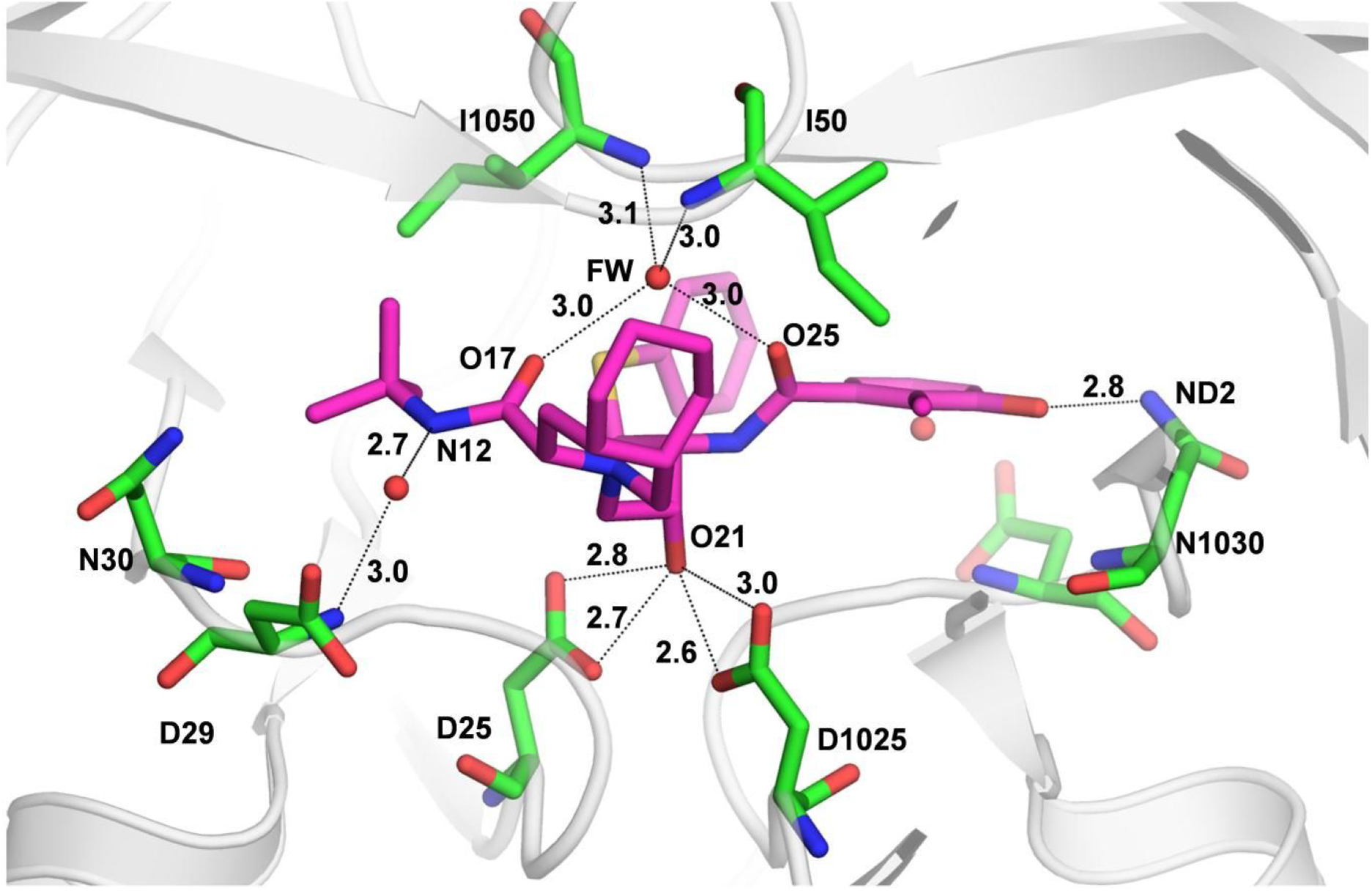
Hydrogen bonding interactions of nelfinavir in D30N HIV-1 protease. Nelfinavir is shown in magenta and protein residues interacting with the drug are shown in green. Flap water is labeled as FW.

### Drug resistance against nelfinavir

Depending on the orientation of the N30 side chain, there can be two different possibilities of hydrogen bonding in the D30N-nelfinavir complex. In case of preferred rotamer, the drug will be an acceptor in an O…H-N hydrogen bond while in the alternative arrangement, where N30 OD1 atom is towards the drug, it will be a donor in an O-H…O hydrogen bond. Our analysis based on rotamer preference, R-factors and hydrogen bonding interactions suggest that both the orientations may be present at different times as they are equally likely. In both the situations, the nature of hydrogen bond in the mutant as compared to the wild type HIV-1 protease is different. A charged acceptor-neutral donor hydrogen bond interaction in wild type HIV-1 protease is changed to a neutral donor-neutral acceptor interaction in D30N HIV-1 protease. The free energy change associated with the ionic hydrogen bond of first condition is estimated to be 16-23 kcal/mol which will be greatly reduced to 3-6 kcal/mol in the alternative condition of mutant protease [Rose, 1993]. Therefore, the change in the nature of hydrogen bond is likely to result into a relatively weaker hydrogen bond between nelfinavir 2-methyl-3-hydroxy benzamide group and amide side chain of D30N HIV-1 protease. Overall conformation of nelfinavir in the mutant structure is also slightly altered from the wild type with all the four sub-sites being affected (Figure 4). These small changes in the conformation of the drug leads to suboptimal fitting of the drug in the active site of mutant protease. Overall normalized complementarity between nelfinavir and active site calculated using LPC software [Sobolev et al 1999], is reduced from 0.79 in the wild type protease to 0.65 in the D30N HIV-1 protease. The reduced complementarity of nelfinavir in D30N HIV-1 protease confirms suboptimal binding of the drug. The decrease in the strength of a crucial hydrogen bond and suboptimal binding of the nelfinavir in the D30N HIV-1 protease active site should result into loss of binding affinity of nelfinavir, leading to the drug resistance observed in HIV/AIDS patients.

### D30N mutation does not provide cross resistance to other drugs

D30N mutation is exclusive to nelfinavir and does not show cross resistance to other drugs except minor resistance against atazanavir and darunavir [Rhee et al 2003; Kovalevski et al 2006]. Interactions of different drugs with the residue 30 observed in different crystal structures are listed in table 2. Nelfinavir makes a direct hydrogen bond with the side chain of the D30 and correspondingly has drastically reduced affinity towards D30N mutant. Ritonavir, indinavir, saquinavir, amprenavir, tipranavir and lopinavir do not form any hydrogen bond with the D30 side chain and their affinities are not affected by D30N substitution. Atazanavir forms a water-mediated hydrogen bond and accordingly, D30N substitution only provides 2.4 ~fold resistance against it [Rhee et al 2003]. Similarly, Kovalevsky et al [2006] have attributed loss in binding affinity of D30N mutant towards TMC-114 (darunavir) to substitution of a direct hydrogen bond by a water-mediated hydrogen bond. Thus, efficacy of D30N as a drug resistance mutation and direct hydrogen bonding of the drug with D30 appears to be correlated.

**Table 2:**
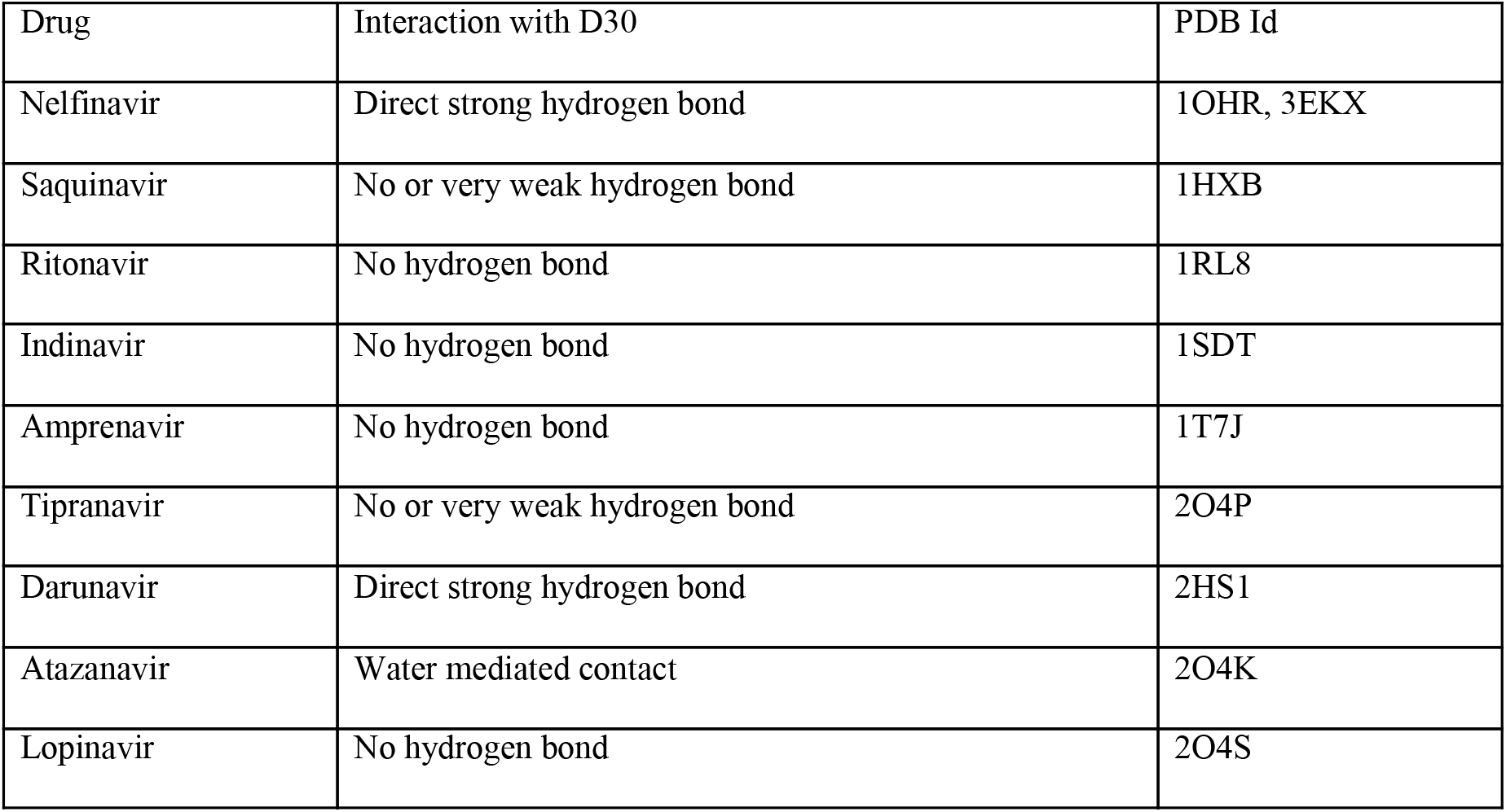
Interaction between different protease inhibitors and side chain of the residue 30 of HIV-1 protease

| Drug | Interaction with D30 | PDB Id |
| --- | --- | --- |
| Nelfinavir | Direct strong hydrogen bond | 1OHR, 3EKX |
| Saquinavir | No or very weak hydrogen bond | 1HXB |
| Ritonavir | No hydrogen bond | 1RL8 |
| Indinavir | No hydrogen bond | 1SDT |
| Amprenavir | No hydrogen bond | 1T7J |
| Tipranavir | No or very weak hydrogen bond | 2O4P |
| Darunavir | Direct strong hydrogen bond | 2HS1 |
| Atazanavir | Water mediated contact | 2O4K |
| Lopinavir | No hydrogen bond | 2O4S |

### D30N reduces cleavage activity of protease and affects replicative capacity of the virus

Mutation D30N leads to substantially reduced cleavage activity of HIV-1 protease and also affects replicative capacity/vitality of the virus. D30 is located in the HIV-1 protease active site and interacts with the peptide substrates through both main chain and side chain atoms [Prabhu-Jeyabalan et al 2002]. The mutation D30N will have two different effects on these interactions: first, removal of charge on the 30^th^ residue will alter the electrostatic interactions with the substrates and is likely to affect the cleavage efficiency of HIV-1 protease with respect to those substrates where this charge interaction plays important role. Second, deviation in main chain and side chain atoms seen in D30N unliganded structure may also effect the hydrogen bonding distance of this interaction and therefore affect those substrates where the strength of this hydrogen bond is important to hold substrates in optimum position and orientation. Prabu-Jeyabalan et al., has determined crystal structures of 6 different substrates in complex with the inactive HIV-1 protease [Prabu-Jeyabalan et al. 2002]. Among these 6 substrates, 4 substrates make direct hydrogen bonds with the D30 side chain. In substrate MA-CA and CA-P2, serine OG atom and glutamate OE1 forms strong hydrogen bonds (2.4-2.5 Å) with Asp30 OD2 while in P2-NC and P1-P6 substrates, Glutamine NE2 interacts with Asp30 side chain (2.9-3.0 Å). In P1-P6, Arg NE also forms hydrogen bonds with Asp30 side chains (3.0-3.2 Å). Mutation D30N is likely to affect these hydrogen bonds and in-turn cleavage process of these substrates will be altered. Any alterations in the cleavage rates will have a large impact on overall fitness of HIV-1 virus. Only three residues D29, D30 and R8 are involved in the side chain hydrogen bonds (less than 3.2Å) with the substrates. Interestingly, all the three residues are absolutely conserved except D30 which mutates under extreme selection pressure of selected drugs particularly nelfinavir. In fact, substrate co-evolution has been observed in HIV-1 protease p1-p6 substrate to compensate for the loss of electrostatic and and hydrogen bonds interactions due to D30N mutation [Kolli et al., 2014]. Thus, overall these altered electrostatic and hydrogen bond interactions are likely to affect the catalytic activity of the enzyme. Residue D30 is part of a structurally conserved loop (24-34) shown to be the part of HIV-1 protease folding core [Wallqvist et al 1998, Broglia et al 2008, Verkhivker et al 2008, Bonomi et al 2010]. Mutation D30N might affect the stability of this folding core, thereby prompting the unfolding of mutant protease at lower temperature. Thus, the effect of D30N mutation on HIV-1 protease folding along with the reduced catalytic activity are likely to be the reason for reduced fitness of the virus carrying D30N HIV-1 protease.

## Conclusion

D30N mutation in HIV-1 protease observed under selective pressure of the drug nelfinavir reduces the binding affinity of the drug nelfinavir and also affects the catalytic process of the enzyme. The structure of mutant protease in complex with nelfinavir shows that a crucial hydrogen bond between the 2-methyl-3-hydroxy benzamide group from the drug and side chain atoms of 30^th^ residue is altered. The nature of this hydrogen bond is changed from ionic O-H…O hydrogen bond to an O-H…N/ O…H-N hydrogen bond in the D30N mutant. The loss of coulombic interaction part of the original interaction in the mutant-drug complex is likely to reduce the strength of the hydrogen bond and therefore results into reduced affinity and drug resistance towards nelfinavir. The structure of D30N HIV-1 protease in unliganded form suggests structural effects that might be responsible for reduced catalytic activity of D30N HIV-1 protease. Since D30 directly interacts with many of the substrates, this mutation also affects substrate binding leading to reduced cleavage activity of the D30N HIV-1 protease. This decreased catalytic activity and reduced stability of folding core due to D30N mutation is suggested as the reason behind reduced replicative capacity of the HIV harboring D30N HIV-1 protease.

## Acknowledgement

We thank late Dr Jean - luc Ferrer, FIP beamline, ESRF, for help in data collection and processing. We also thank the National Facility for Structural Biology for providing the facility to conduct these experiments.

